# The haplotype-resolved assembly of COL40 a cassava (*Manihot esculenta*) line with broad-spectrum resistance against viruses causing Cassava brown streak disease unveils a region of highly repeated elements on chromosome 12

**DOI:** 10.1101/2024.09.30.615795

**Authors:** Corinna Thoben, Boas Pucker, Stephan Winter, Bethany Fallon Econopouly, Samar Sheat

## Abstract

Cassava (*Manihot esculenta Grantz*) is a vital staple crop for millions of people, particularly in Sub-Saharan Africa, where it is a primary source of food and income. However, cassava production is threatened by several viral diseases, including cassava brown streak disease, which causes severe damage to the edible storage roots. Current cassava varieties in Africa lack effective resistance to this disease, leading to significant crop losses. We investigated the genetic diversity of cassava and identifed new sources of resistance to the viruses causing cassava brown streak disease. The cassava line, COL40, from a South American germplasm collection showed broad-spectrum resistance against all known strains of the viruses that cause this disease. To further understand the genetic basis of this resistance, we sequenced the genome of COL40 and produced a high-quality, haplotype-resolved genome assembly. This genomic resource provides new insights into cassava’s genetic architecture, particularly in regions associated with disease resistance. The sequence reveals significant structural variation, including transposable elements, inversions, and deletions, which may contribute to the resistance phenotype. The reference genome assembly presented here will provide a valuable genomic resource for studying the cassava brown streak resistance and will help in accelerating breeding efforts to introduce virus resistance into African cassava varieties. By identifying genetic variants linked to resistance, future breeding programs can develop cassava cultivars that are more resilient to viral threats, enhancing food security and livelihoods for smallholder farmers across regions affected by the disease.

## Introduction

Cassava (*Manihot esculenta*) is an important world food crop that produces storage roots rich in starch to provide a main source of food and income for people in Africa, South America, India, and Southeast Asia. The plant originates from the Amazon region of South America but is especially important in Sub-Saharan Africa, where half of the world’s cassava is produced mainly by smallholder farmers. This resilient plant adapts well to diverse environments, is resistant to drought, grows in poor soils and guarantees food security even in the most adverse environmental conditions. Nevertheless, cassava cultivation is constrained by pests and diseases, and because stem cuttings are taken to propagate the crop, diseases are maintained and disseminated by vegetative propagules. Viruses are associated with cassava cultivation in all cassava regions of the world and present serious threats to crop production. In Africa, viruses causing cassava mosaic disease are endemic, but the deployment of varieties with high virus resistance has mitigated the impact of the diseases. In contrast, there was no means to control the spread of the viruses causing cassava brown streak disease (CBSD), which has a severe impact because of the root necrosis destroying the edible tubers of the crop The relatively recent outbreak, first noted in 2004/2005 in Uganda (Alicai *et al*., 2007), the limited knowledge about the etiology and epidemiology of the disease, and the lack of virus-resistant cassava in Africa delayed comprehensive actions to control the disease. Searching for new sources of resistance, we explored the diversity of South American cassava. We infected a subset of a germplasm collection held at CIAT with diverse cassava brown streak viruses to identify lines that showed resistance to either isolates of the species cassava brown streak virus (CBSV) or against the Uganda cassava brown streak virus (UCBSV). After stringent infection and virus screening, three germplasm accessions COL2182, PER556, and COL40 showed broad-spectrum immunity to all species and strains of the viruses causing CBSD (Sheat *et al*., 2019). COL40, the most promising line, is now used as a source to provide CBSD resistance essentially in all cassava improvement programs. First prototypes from crosses with African lines are in the field, proving that high virus resistance in progenies from COL40 can be reached. Progress has been made in characterizing the resistance phenotype of COL40 (Sheat, Margaria & Winter, 2021) and advancing field evaluation and resistance assessment (Sheat & Winter, 2023). However, important steps towards molecular breeding, genomic prediction, marker-assisted selection and a mechanistic explanation of the virus resistance in COL40 remain open challenges.

The reconstruction of complete, haplotype-resolved genome sequences of cassava genotypes is a substantial achievement in providing genome information and resources to advance our understanding of the cassava genome(s) and accelerate breeding to future-proof the crop for resilience to biotic and abiotic stresses, agronomic traits and consumer demands. As a highly heterozygous species (Prochnik *et al*., 2012) (1% in TME 204; Qi *et al*., 2022) its genome comprises extensive amounts of repetitive elements (TMEB117 >60% ; (Landi *et al*., 2023), allelic variation, inversions, and deletions/ insertions comprising large segments of the genome (Ramu *et al*., 2017). This genomic structural variation limits the use of a single reference genome sequence (AM560-2; Phytozome 13, v 8.1) that does not capture the full complement of sequence diversity of a crop species (Jayakodi *et al*., 2020) and may lack genes present in other genotypes, e.g. NBS-LRR genes relevant for disease resistance (Lozano *et al*., 2015).

Recently, high-quality haplotype-resolved genome sequences of African cassava lines were reported (Qi *et al*., 2022; Kuon *et al*., 2019; Mansfeld *et al*., 2021; Landi *et al*., 2023). The lines chosen were TME3 and 60444, to contrast a landrace with high resistance against viruses causing CMD, carrying the CMD2 resistance locus (TME3) and a highly susceptible line 60444. Similarly, TME204 (Qi *et al*., 2022) carrying CMD2 and TMEB117 lacking (Landi *et al*., 2023) were chosen to provide high-resolution genome data for virus studies on resistance mechanisms and, to guide future breeding. Our study follows a similar rationale. By generating a high-quality COL40 genome sequence we now have a reference to center our research, to advance our understanding about virus resistance/ immunity in cassava, identify genetic variants associated with resistance traits (GWAS) and to inform cassava resistance breeding using COL40 as source.

## Material & Methods

### Plant cultivation

Cassava COL40 plantlets grown in tissue culture were transferred to pots and maintained at the greenhouse facility of the DSMZ Plant Virus Department in Braunschweig Germany at temperatures between 26 °C to 32 °C and >75% relative humidity with additional light provided during German winter conditions.

### DNA Extraction, library preparation and sequencing

Prior to sampling, the plants were transferred to the dark for 24 hours after which fully-expanded leaves (between 3^rd^ to 5^th^ leaf) were taken and flash frozen in liquid N_2_. Leaves (10-15 g) were sent to Arizona Genomics Institute (AGI) for high-molecular-weight (HMW) genomic DNA (gDNA) extraction, library preparation, and sequencing. HMW gDNA extraction was done essentially following the CTAB method of Doyle and Doyle,1987 (Doyle & Doyle, 1987) with minor modifications (Fu, Song & Jameson, 2017). The HiFi express template prep kit 2.0 was used for PacBio HiFi SMRTbell library construction. Fragments between 10 to 25-kb were selected for HiFi sequencing on the Sequel II instrument. SeqII v.2.0 chemistry was used with two SMRT cells 8M v1 in circular consensus sequencing (CCS) mode for a 30-hour run time per cell.

For RNAseq analysis, leaf, stem and root samples were taken from COL40 and flash frozen in liquid N_2_ prior to extraction of total RNA using a RNA extraction kit following the manufacturer’s protocol (Epoch, United States). RNA was quantified in a QubitR fluorometer (Thermo Fisher Scientific, United States) using the Qubit RNA BR Assay Kit (Thermo Fisher Scientific, United States), and checked for size distribution in a bioanalyzer (Agilent) prior to sequencing at Novogene (Science Park, Milton, Cambridge, UK). At Novogene, mRNA was purified from total RNA preparations using poly-T magnetic beads, fragmented and subjected to first strand cDNA synthesis using random hexamer primers followed by second strand cDNA synthesis and end repair. Library construction, quality controls, and paired-end sequencing (PE 150) on an Illumina Novaseq 6000 platform followed Novogene’s mRNA sequencing workflow.

### Assembly generation and quality control

The HiFi reads of the COL40 cultivar were assembled by Hifiasm 0.19.8-r603 (Cheng *et al*., 2021; Cheng *et al*., 2022), NextDenovo 2.5.0 (https://doi.org/10.1186/s13059-024-03252-4), Flye 2.9.3-b1797 (Kolmogorov *et al*., 2019), and Shasta 0.11.1 (Shafin *et al*., 2020). To avoid compatibility issues with tools in downstream analysis, contig identifiers of the assemblies were cleaned using the python script clean_genomic_fasta.py v0.15 (Meckoni, Nass & Pucker, 2023). Assembly statistics were calculated with the python script contig_stats.py v1.31 (Pucker *et al*., 2016) Completeness of universal single-copy orthologs was checked with BUSCO 5.2.2 and the eudicots_odb10 dataset (Manni *et al*., 2021). A k-mer based analysis was performed with Merqury (Rhie *et al*., 2020) and k-mers generated from the cultivars HiFi reads (k=21). The assembly’s coverage was analyzed by mapping the HiFi reads to the assembly with minimap2 2.24-r1122 and “-ax map-hifi –secondary=no” option (Li, 2018). Samtools 1.17-29-gcc18465 (Bonfield *et al*., 2021) was applied for conversion and sorting of the mapping file. The coverage was calculated with the “genomecov” command of bedtools v2.30.0 (Quinlan & Hall, 2010).

### Scaffolding of the assembly

Genetic markers of the composite genetic map of *Manihot esculenta Crantz* by the International Cassava Genetic Map Consortium (ICGMC) (File S2,(Consortium, 2015)) were used for scaffolding. The genetic markers were mapped to the assemblies using the python script genetic_map_to_fasta.py v0.2 (https://github.com/c-thoben/CassavaGenomicsProject), which creates a csv map based on the best BLAST hits for each marker with minimum similarity of 99.0 % and score of 175. The scaffolding was performed by converting the csv map with ALLMAPS (JCVI utility libraries 1.4.2) (Tang *et al*., 2015)) “merge” option and path construction using the “path” option. Scaffold statistics were calculated with the python script contig_stats.py v1.31 (Pucker *et al*., 2016).For the Hifiasm assembly, the scaffolding was performed for each haplophase separately. The scaffolds of both haplophases were merged together and for each scaffold, a suffix was added to the scaffold IDs indicating its haplophase. The coverage plot of the chromosomes including the density of the transposable elements was created with the python script chromosome_coverage_te_plot.py v0.4 (Pucker et al., 2016), (https://github.com/c-thoben/CassavaGenomicsProject).

### Prediction and functional annotation of polypeptide sequences

In total, 21 RNA-Seq libraries of the COL40 cultivar (ERS20926298, ERS20926299, ERS20926300, ERS20926301, ERS20926302, ERS20926303, ERS20926304, ERS20926305, ERS20926306, ERS20926307, ERS20926308, ERS20926309, ERS20926310, ERS20926311, ERS20926312, ERS20926313, ERS20926314, ERS20926315, ERS20926316, ERS20926317, ERS20926318) and 24 further RNA-Seq libraries (SRR19351399, SRR19351400, SRR19351401, SRR19351402, SRR25288231, SRR25288232, SRR25288235, SRR25288236, SRR25537338, SRR25537339, SRR25537340, SRR3629818, SRR3629824, SRR3629835, SRR3629837, SRR3629838, SRR3629840, SRR3629842, SRR3629843, SRR3629845, SRR3629850, SRR3629851, SRR3629853, SRR3629859) from other cultivars were mapped to the COL40 assembly with HISAT2 2.2.1(Zhang *et al*., 2021), (Kim *et al*., 2019), (Kim, Langmead & Salzberg, 2015; Pertea *et al*., 2016) and “--dta” option. Samtools 1.17-29-gcc18465 (Danecek *et al*., 2021) was applied for conversion and sorting of the mapping files (Bonfield *et al*., 2021).Together with external protein hints from *Manihot esculenta* Crantz v8.1 (Consortium, 2015) and the OrthoDB 11 Viridiplantae database (Kuznetsov *et al*., 2022) the mapped RNA-Seq reads were given to BRAKER3v3.0.6 (Stanke *et al*., 2008; Stanke *et al*., 2006; Gabriel *et al*., 2021; Gabriel *et al*., 2024; Brůna, Lomsadze & Borodovsky, 2024; Kovaka *et al*., 2019; Pertea & Pertea, 2020; Quinlan, 2014; Brůna, Lomsadze & Borodovsky, 2020; Buchfink, Xie & Huson, 2015; Lomsadze et al., 2005; Iwata & Gotoh, 2012; Gotoh, Morita & Nelson, 2014; Li et al., 2009; Barnett et al., 2011) as external hints to perform the structural annotation. Based on the mapped RNA-Seq reads, the coverage of the structural annotation was analyzed with the python script RNAseq_cov_analysis.py v0.1 (Meckoni *et al*., 2023), https://github.com/bpucker/GenomeAssembly/) and filtered for transcripts with a coverage > 90 %. Completeness of the predicted polypeptide sequences was checked with BUSCO v5.2.2 and the eudicots_odb10 (Manni *et al*., 2021) dataset for each haplophase separately. The polypeptide sequences were functionally annotated using InterProScan5 v5.67-99.0 (Manni *et al*., 2021; Blum *et al*., 2021).

### Prediction and analysis of transposable elements

Transposable elements were predicted with EDTA v2.1.0 (Ou *et al*., 2019; Ellinghaus, Kurtz & Willhoeft, 2008; Xu & Wang, 2007; Ou & Jiang, 2019), (Ou & Jiang, 2018), (Su, Gu & Peterson, 2019; Shi & Liang, 2019; Xiong *et al*., 2014; Zhang *et al*., 2022) and the “–overwrite 1 –sensitive 1 -anno 1 -evaluate 1” options. Annotated coding sequences predicted by BRAKER3 were provided with the “--cds” option. Annotated transposable elements were further analyzed with the script COL40_TE_repeat_analysis.ipynb (https://github.com/c-thoben/CassavaGenomicsProject) based on the R script TMEB117TEandGeneAnnotation.R provided by (Landi *et al*., 2023), (https://github.com/LandiMi2/GenomeAssemblyTMEB117). The circlize package (Gu *et al*., 2014) was used to calculate the density of transposable elements and predicted coding sequences and visualize them in a circos plot created with the R package circlize (Gu *et al*., 2014). The coverage plot of the chromosomes including the mean coverage over 1 kbp and the density of the transposable elements over 1 Mbp was created with the python script coverage_te_plot.py v0.5, (Pucker *et al*., 2016), (https://github.com/c-thoben/CassavaGenomicsProject).

## Results

### Assembler comparison and quality control

The HiFi dataset consisted of 3,440,114 reads with an N50 length of 18,106 and 38 % GC content. All assemblies were > 80 % complete representations of the genome according to the k-mer analysis and, with exception of the Shasta assembly, show a high completeness regarding the single-copy universal orthologs. The Hifiasm (N50: 35 Mbp, 30 Mbp) and NextDenovo (N50: 6,7 Mbp) assemblers performed best in producing assemblies with a high continuity (Table 1).

**Table 1.** Assembly statistics and quality control of assemblies

|  | Hifiasm | NextDenovo | Flye | Shasta |
| --- | --- | --- | --- | --- |
| **Assembly size** | 823 MB, 694 Mbp | 804 Mbp | 693 Mbp | 1015 Mbp |
| **Number of contigs** | 2605, 446 | 394 | 9217 | 7692 |
| **N50 of contigs** | 35 MB, 30 Mbp | 6,7 Mbp | 119 kbp | 1,6 Mbp |
| **Number of scaffolds** | 2600, 427 | 224 | 6976 | 6334 |
| **N50 of scaffolds** | 37 MB, 36 Mbp | 41 Mbp | 545 kbp | 41 Mbp |
| **BUSCO** | 98.8 % (D: 8.6 %), 98.8 % (D: 8.4 %) | 99.0 % (D: 13.4 %) | 98.8 % (D: 17.2 %) | 75.5 % (D: 58.9 %) |
| **K-mer completeness** | 82, 81 (asm: 98) | 86 | 86 | 82 |
| **QV** | 57, 63 (asm: 58) | 50 | 36 | 61 |

The Hifiasm assembly had the highest k-mer completeness (98 %, ∼80 % for each haplophase) and a good assembly consensus quality value (QV) of 58 according to the k-mer analysis. The analysis of the k-mer copy numbers (Fig. 1) demonstrates that the haplotypes are well-resolved in this assembly.

**Fig. 1.**
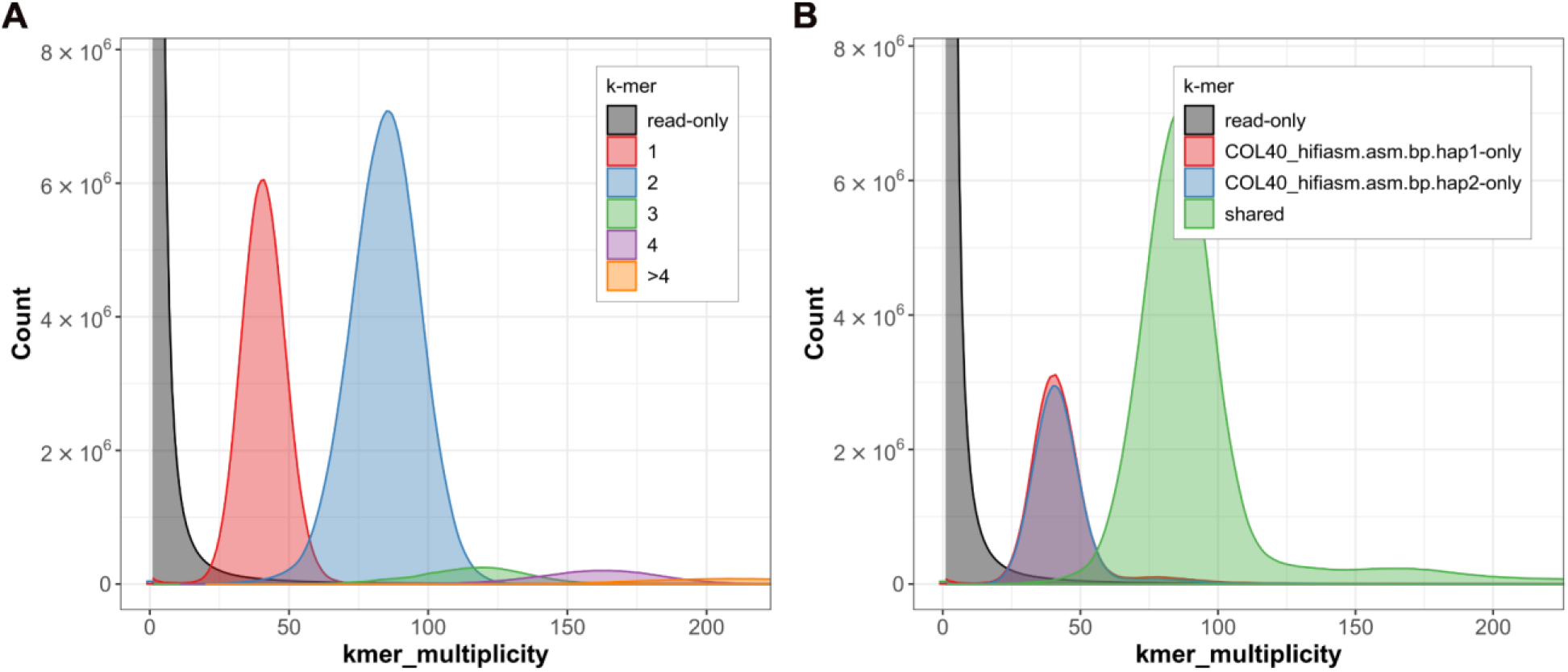
Assembly completeness according to k-mer analysis of the assembly. (A) Merqury copy number spectrum plot of the HiFi reads. The red peak represents 1-copy reads. The blue peak represents 2-copy reads. HiFi reads not included in the assembly are represented by the gray peak. (B) Copy number spectrum plot of the k-mers in A. Heterozygous reads are colored red or blue according to the haplophase. Homozygous reads or haplotype-specific duplications are colored green.

The Shasta assembly had the highest assembly quality (QV 61) in the k-mer analysis, standing out in this specific criterion, but was outperformed by the Hifiasm assembly in the remaining criteria. Therefore, the Hifiasm assembly was chosen as the representative genome sequence of COL40.

### Scaffolding and structural annotation

In the scaffolding process, 16,233 out of 22,403 markers were mapped to the contigs of haplophase A and used to scaffold 23 contigs representing the 18 *M. esculenta* chromosomes (Additional File 1, Additional File 2). The 18 chromosomes of haplophase B were scaffolded from 37 contigs with reference to 16,169 mapped markers. The remaining 2,582 contigs for haplophase A and 409 contigs for haplophase B remained unplaced. The structural annotation of the Hifiasm assembly predicted 82,151 polypeptide sequences for haplophase A and 72,383 polypeptide sequences for haplophase B. After filtering the transcripts based on their coverage in the mapped RNA-Seq reads, the number of sequences was reduced to 36,064for haplophase A and 34,029for haplophase B. The BUSCO analysis revealed a high completeness of 96.6 % (D: 22.7 %) for haplophase A, 96.7 % (D: 21.7 %) for haplophase B, and 98.6 % (D: 94.6 %) for both haplophases.

### Assembly coverage analysis

The coverage analysis revealed an average coverage of 40.4-fold for haplophase A and 39.7-fold for haplophase B. The coverage plot demonstrates an equal distribution of the coverage, which can be attributed to the high resolution of the haplophases in the assembly (Additional File 3, Additional File 4). For both haplophases, a coverage drop at the end of chromosome 12 (downstream of ∼ 39 Mbp for haplophase A, ∼ 37 Mbp for haplophase B) is displayed in the coverage plot (Fig. 2). This is also reflected in the coverage histogram of chromosome 12, which shows a second peak around a coverage of 15 for both haplophases (Additional File 3, Additional File 4). For both haplophases, the density of transposable elements is enhanced in these regions, which is further discussed below (Fig. 2 & 3).

**Fig. 2.**
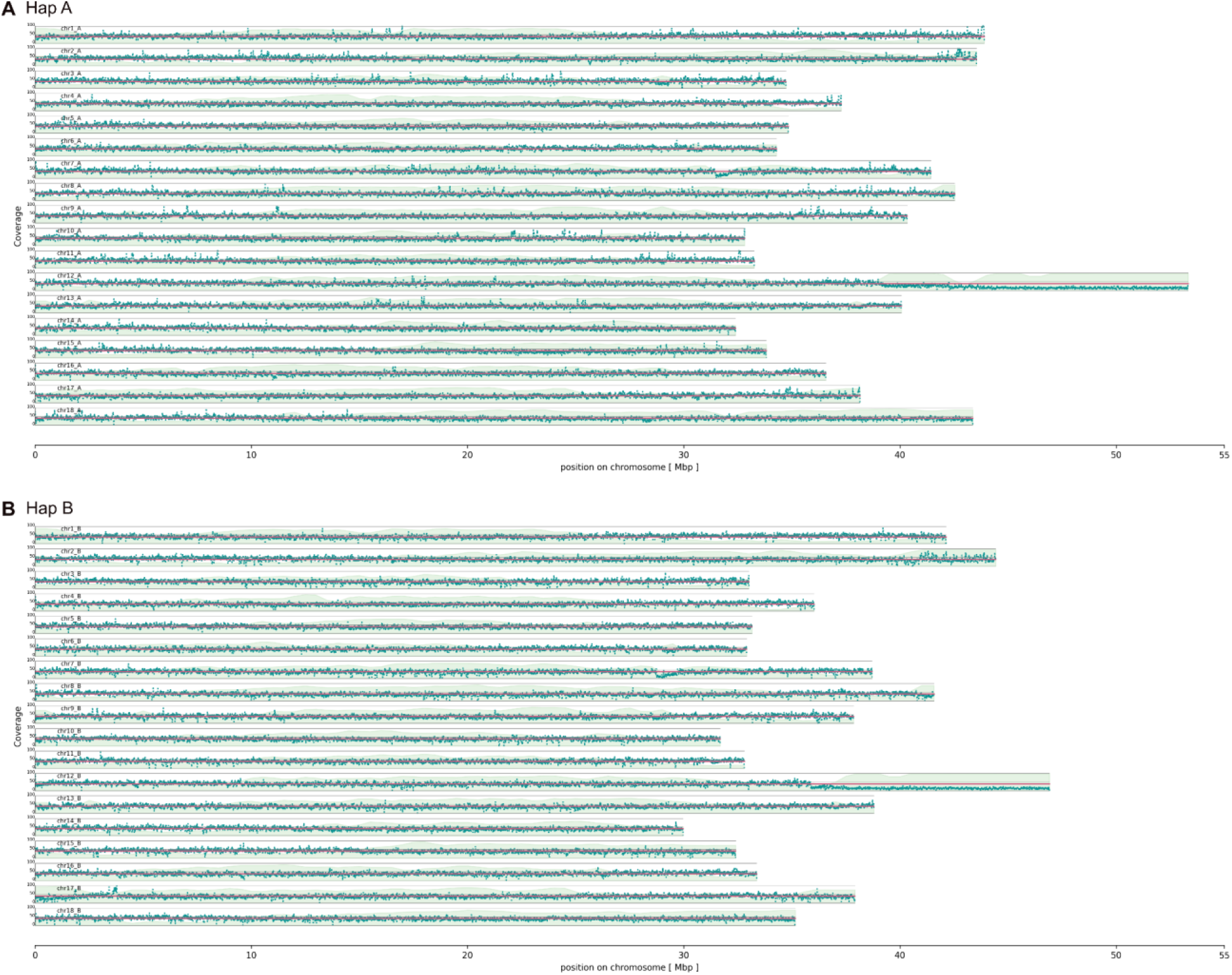
Coverage plot of the chromosome-resolved Hifiasm assembly. The HiFi reads were mapped to the haplophase A (top) and haplophase B (bottom) of the scaffolded assembly and the mean coverage was calculated for each haplophase (red line) and block-wise over 1 kbp intervals for each chromosome (dark green dots). The density of transposable elements was calculated block-wise per1 Mbp interval (light green area).

### Annotation of transposable elements

In total, 3027 repeat regions were annotated, which cover 58.04 % of the COL40 assembly. The most abundant transposable elements (TL) are long terminal repeats – retrotransposons (LTR-RTs), mostly Gypsy LTR-RT, which cover 51.88 % of the complete assembly and are the most abundant family on all chromosomes (Fig. 3).

**Fig. 3.**
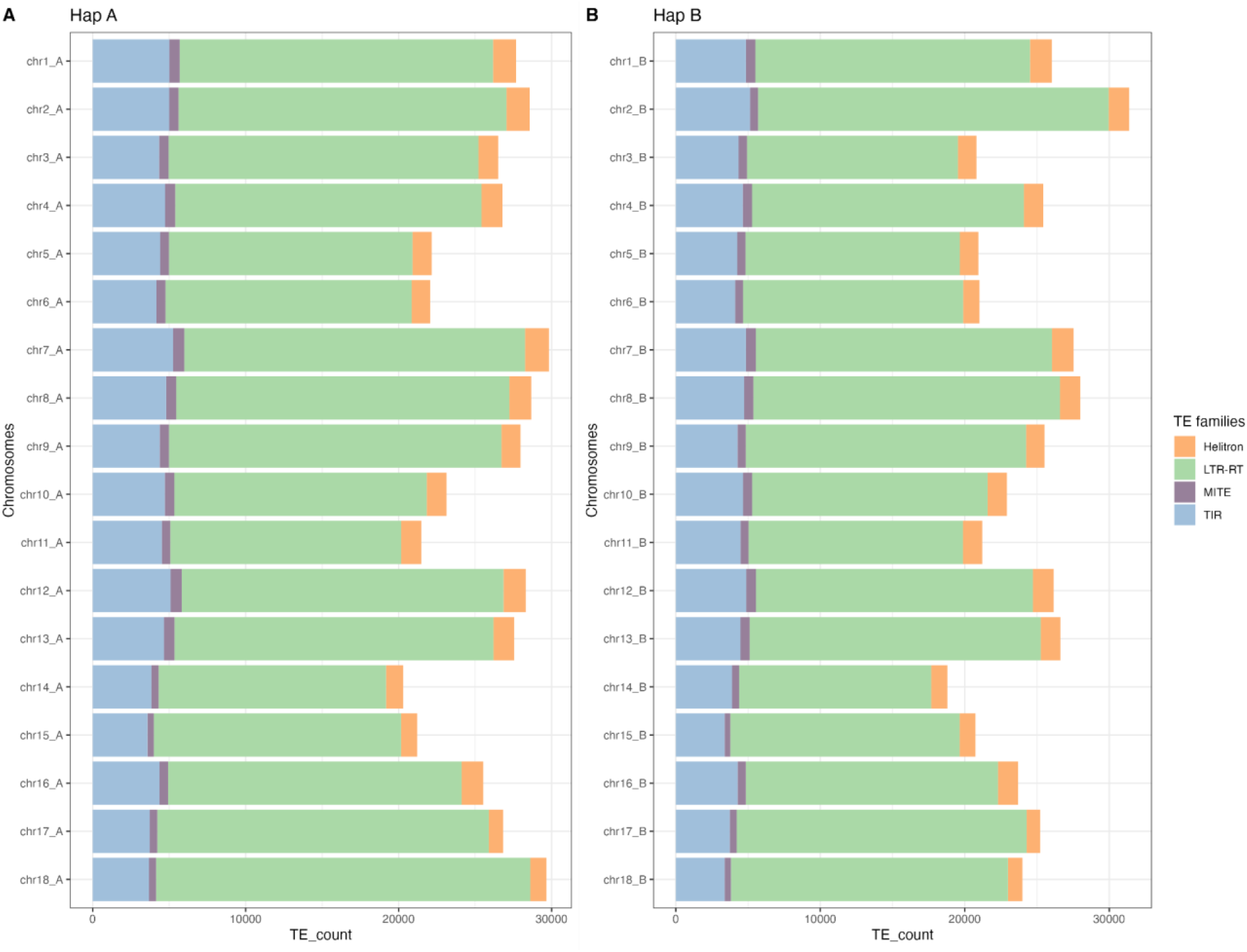
Distribution of transposable element families for pseudochromosomes of haplophase A (left) and haplophase B (right) in the Hifiasm assembly. Unclassified transposable elements were removed from the annotation. For all pseudochromosomes, LTR-RTs are the most abundant repeats for both haplophases.

As described above, the density of TE increases at the end of chromosome 12 in both haplophases (Fig. 4). This increase can be attributed to fragmented transposable elements, as the density of structurally intact transposable elements decreases. Furthermore, the density of predicted coding sequences is distinctly reduced in this region. The scaffolding results (Additional File 1, Additional File 2) show that no markers of the *M. esculenta* Crantz genetic map by the International Cassava Genetic Map Consortium (ICGMC) (File S2,(Consortium, 2015) were mapped to the region, suggesting that the transposable elements in this area are either not represented in the map or that markers could not be developed due to the high level of repetitive sequences present in the region.

**Fig. 4.**
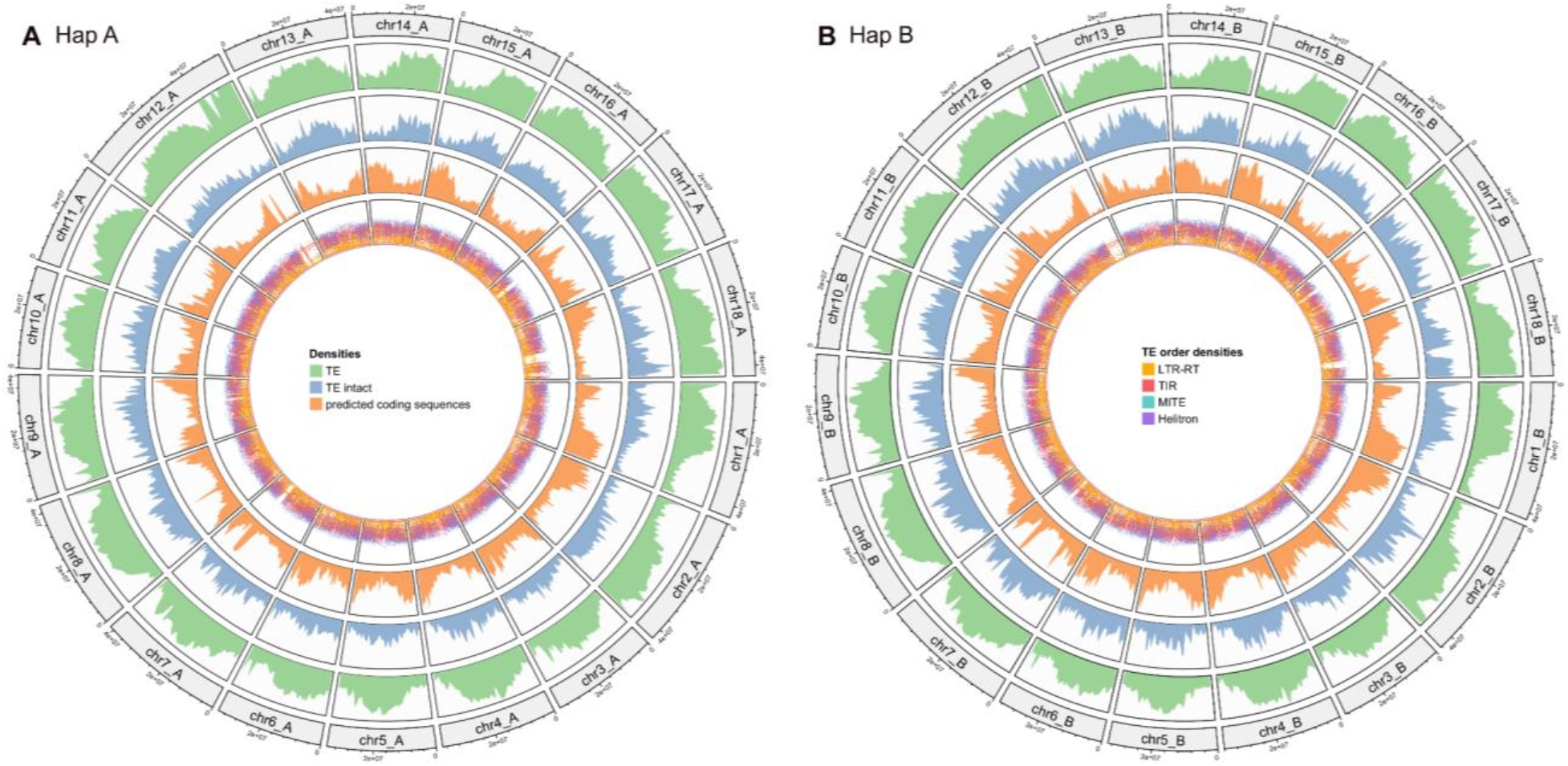
Density of transposable elements and predicted coding sequences across the pseudochromosomes of haplophase A (left) and haplophase B (right) in the Hifiasm assembly. For all tracks, the density was calculated over a window size of 1 Mbp. The density of transposable elements is shown in the outer track and reflects the percent of the window covered by the input regions (green). The second track shows the density of structurally intact transposable elements (blue). The density of predicted coding sequences is displayed in the third track (orange). In the inner track, a rainbow plot represents the minimal distance (log_10_-transformed) to the neighboring regions. The color of the region displays the transposable elements order.

## Discussion

In this study, we presented a haplotype-resolved genome assembly of the cassava line COL40, a South American germplasm line with high and broad-spectrum resistance to the viruses causing CBSD. The assembly of COL40 joins a growing collection of high-quality genome assemblies from other cassava cultivars, such as TMEB117, TME204, and TME3 (Kuon *et al*., 2019; Qi *et al*., 2022; Landi *et al*., 2023) and highlighting the increasing role of advanced sequencing technologies in improving crop genomics.

Using PacBio HiFi reads and after a comparative evaluation of assembly tools, we selected Hifiasm as the optimal assembler due to its superior continuity and completeness. Hifiasm also generated best assembly in other cassava genome studies (Kuon *et al*., 2019; Landi *et al*., 2023) with haplophase sizes between 665 Mbp and 823 Mbp (Table 2).

**Table 2.** Comparison of recently published cassava assemblies

|  | COL40 | TMEB117 | TME204 | TME3 |
| --- | --- | --- | --- | --- |
| **Chromosomes resolved** | yes | yes | yes | No |
| **Haplotype-resolved** | yes | yes | yes | No |
| **Assembly size** | 823 MB, 694 Mbp | 694 MB, 665 Mbp | 762 MB, 706 Mbp | 1225 Mbp |
| **N50 of scaffolds** | 37 MB, 36 Mbp | 38 MB, 36 Mbp | 18 MB, 26 Mbp | 53 Mbp |
| **BUSCO** | 98.8 % (D: 8.6 %), 98.8 % (D: 8.4 %) | 98.9 % (D: 8.8 %), 98.9 % (D: 8.8 %) | 99 % (D: 4.9 %), 98.8 % (D: 4.2 %) | 94.8 % (D: 19.7 %) |
| **K-mer completeness** | 82 %, 81 % (asm: 98 %) | 78.63 %, 77.95 % (asm: 98.79 %) | 79.6 %, 79.1 % (asm: 98.4 %) | - |
| **QV** | 57, 63 | 64, 68 | 45, 49 | - |
| **BUSCO (Predicted polypeptide sequences)** | 96.6 % (D: 22.7 %), 96.7 % (D: 21.7 %) | ~ 90 % both | 96.7 % (D: 17.4 %), 96.7 % (D: 16 %) | - |
| **TE proportion** | 58.04 % | 57.37 %, 54.42 % | > 60 % | 64.81 % |
| **Reference** | This study | Landi *et al.*, 2023<br>https://doi.org/10.1038/s41597-023-02800-0 | Kuon *et al.*, 2019<br>https://doi.org/10.1186/s12915-019-0697-6 | Qi *et al.*, 2022<br>https://doi.org/10.1093/gigascience/gi-ac028 |

The high accuracy of HiFi data was crucial for achieving a robust genome assembly with phased haplotypes. Similarly, PacBio HiFi sequencing reported for the assembly of other plant genomes, including *Haloxylon ammodendron* (Wang *et al*., 2022), and *Rhododendron vialii* (Chang *et al*., 2023) allowed generation of highly contiguous genome assemblies for diploid and polyploid species and made it ideal for resolving the complex structure of cassava’s genome.

All assemblies present a high completeness of > 98 % in both BUSCO and k-mer analysis and around 80 % for each haplophase in k-mer analysis (Table 2). Our results are consistent with other assemblies of complex plant genomes like that of the European common bean (Carrère *et al*., 2023). This level of completeness indicates that our assembly has captured the vast majority of the gene space, an essential feature for downstream functional studies, particularly for understanding CBSD resistance mechanisms.

Comparing the QV of different cassava assemblies (Table 2), we observed that the COL40 genome assembly had a QV of 57 for one haplotype and 63 for the other, which is close to the TMEB117 assembly (Landi *et al*., 2023). In contrast, earlier assemblies, such as TME204 (Qi *et al*., 2022) had significantly lower QVs. These improvements in genome quality metrics reflect the advancements made by HiFi sequencing, as highlighted by Sun *et al*., (2022), where HiFi reads produced near-perfect assemblies for various complex plant genomes.

Another outstanding feature of our assembly is that we resolved high-quality polypeptide sequences, with a completeness of around 97% for both haplotypes. These results align with other high-quality plant genome assemblies produced using HiFi data, such as the assembly of *Salix wilsonii* (Han *et al*., 2022) and the bean genome (Carrère *et al*., 2023). These findings underscore the value of using HiFi reads, as they produce long, highly accurate reads that are crucial for resolving complex regions and providing reliable gene annotations.

The COL 40 assembly provids critical insights into the repetitive regions and structural variations in cassava genomes. The coverage analysis along the pseudochromosomes after scaffolding of the COL40 assembly revealed a region with reduced coverage at the end of chromosome 12 on both haplophases (Fig. 2). The annotation of this assembly region showed a high density of fragmented transposable elements TE (Fig. 4) which was also found in chromosome 12 of TMEB117 (Landi *et al*., 2023). This region is not covered by genetic markers in the composite genetic map of *M. esculenta* Crantz by the International Cassava Genetic Map Consortium (ICGMC) (File S2), suggesting that the TEs in this region are either not represented or, that the high frequency of repetitive sequences hindered the developed of respective markers. Nevertheless, this region may not be present in all cassava genomes but it is also likely that earlier assemblies based on short sequencing reads may have not resolved repetitive regions and consequently created gaps and fragmentation of the transposable elements. Highly TE-rich regions present challenges to the completeness of genome assemblies, for cassava and other species (Benham *et al*., 2024).

The COL40 assembly focuses on haplotype resolution and genome completeness. In contrast, the telomere-to-telomere assembly of the cassava Xinxuan 048 delved deeper into epigenetic analysis (Xu *et al*., 2023) highlighting that the structural variation between the haploid genomes of Xinxuan 048 was mainly due to TE insertions. The variation led to differences in CG methylation of alleles, thus resulting in differential allelic expression. These findings suggest that differential gene expression in Xinxuan 048 could be influenced by methylation changes caused by varying TE insertions, The proportion of TE elements in the genome of COL 40 is around 60 %, and this is consistently found in all cassava genome assemblies (Kuon *et al*., 2019; Landi *et al*., 2023). These TE-rich regions may hold essential genes for interesting traits, such as those associated with CBSD resistance.

In conclusion, the high-quality, haplotype-resolved genome assembly of cassava COL40 contributes significantly to cassava genomic resources. It lays the groundwork for further comparative analyses between cassava cultivars and provides an invaluable resource to advance our understanding of virus resistance and immunity.

## Availability of data

The data that support the findings of this study have been deposited into:

- HiFi reads of the COL40 cultivar are available from the European Nucleotide Archive (ENA): ERS20926319
- The 21 COL40 RNA-Seq data sets are also available via ENA: ERS20926298, ERS20926299, ERS20926300, ERS20926301, ERS20926302, ERS20926303, ERS20926304, ERS20926305, ERS20926306, ERS20926307, ERS20926308, ERS20926309, ERS20926310, ERS20926311, ERS20926312, ERS20926313, ERS20926314, ERS20926315, ERS20926316, ERS20926317, ERS20926318, and ERS20926319 The assembly and the corresponding annotation are published in LeoPARD: https://leopard.tu-braunschweig.de/receive/dbbs_mods_00077998
- Scripts used for the generation and analysis of the genome sequence assembly are available via GitHub repository: https://github.com/c-thoben/CassavaGenomicsProject

## Acknowledgments

We are grateful for all assistance in the experiments provided by Marianne Koerbler, Michelle “Skadi” Falk and Agnes Pietruszka for the excellent and skillful handling of all cassava plant materials, tissue cultures and nucleic acid preparations. This work was supported by the BMBF-funded de.NBI Cloud within the German Network for Bioinformatics Infrastructure (de.NBI) (031A532B, 031A533A, 031A533B, 031A534A, 031A535A, 031A537A, 031A537B, 031A537C, 031A537D, 031A538A).

## Author contributions

S.S, S.W. and B.P. conceived the idea; C.T. and B.P., devised the bioinformatic approach; S.S. and SW handled the plant materials and RNA-seq; B.F.E. handled PacBio HIFI sequencing; C.T. analyzed the data and compiled the workflows. All authors contributed to the original draft of the manuscript, and reviewed and approved the final manuscript.

## Supplementary Files

- Additional File 1: Scaffolding orientation for each chromosome of haplophase A created with ALLMAPS
- Additional File 2: Scaffolding orientation for each chromosome of haplophase B created with ALLMAPS
- Additional File 3: Coverage histograms for each chromosome of haplophase A
- Additional File 4: Coverage histograms for each chromosome of haplophase B

## Literature cited

Alicai, T., Omongo, C., Maruthi, M., Hillocks, R., Baguma, Y., Kawuki, R., Bua, A., Otim-Nape, G. & Colvin, J. (2007). Re-emergence of cassava brown streak disease in Uganda. Plant Disease 91(1), 24–29.

Barnett, D. W., Garrison, E. K., Quinlan, A. R., Strömberg, M. P. & Marth, G. T. (2011). Bamtools: a C++ API and toolkit for analyzing and managing BAM files. Bioinformatics 27(12), 1691–1692.

Benham, P. M., Cicero, C., Escalona, M., Beraut, E., Fairbairn, C., Marimuthu, M. P. A., Nguyen, O., Sahasrabudhe, R., King, B. L., Thomas, W. K., Kovach, A. I., Nachman, M. W. & Bowie, R. C. K. (2024). Remarkably High Repeat Content in the Genomes of Sparrows: The Importance of Genome Assembly Completeness for Transposable Element Discovery. Genome Biol Evol 16(4).

Blum, M., Chang, H.-Y., Chuguransky, S., Grego, T., Kandasaamy, S., Mitchell, A., Nuka, G., Paysan-Lafosse, T., Qureshi, M. & Raj, S. (2021). The InterPro protein families and domains database: 20 years on. Nucleic Acids Research 49(D1), D344-D354.

Bonfield, J. K., Marshall, J., Danecek, P., Li, H., Ohan, V., Whitwham, A., Keane, T. & Davies, R. M. (2021). HTSlib: C library for reading/writing high-throughput sequencing data. Gigascience 10(2).

Brůna, T., Lomsadze, A. & Borodovsky, M. (2020). GeneMark-EP+: eukaryotic gene prediction with self-training in the space of genes and proteins. NAR genomics and bioinformatics 2(2), lqaa026.

Brůna, T., Lomsadze, A. & Borodovsky, M. (2024). GeneMark-ETP significantly improves the accuracy of automatic annotation of large eukaryotic genomes. Genome Research.

Buchfink, B., Xie, C. & Huson, D. H. (2015). Fast and sensitive protein alignment using DIAMOND. Nature Methods 12(1), 59–60.

Carrère, S., Mayjonade, B., Lalanne, D., Gaillard, S., Verdier, J. & Chen, N. W. (2023). First whole genome assembly and annotation of a European common bean cultivar using PacBio HiFi and Iso-Seq data. Data in Brief 48, 109182.

Chang, Y., Zhang, R., Ma, Y. & Sun, W. (2023). A haplotype-resolved genome assembly of Rhododendron vialii based on PacBio HiFi reads and Hi-C data. Scientific Data 10(1), 451.

Cheng, H., Concepcion, G. T., Feng, X., Zhang, H. & Li, H. (2021). Haplotype-resolved de novo assembly using phased assembly graphs with hifiasm. Nature Methods 18(2), 170–175.

Cheng, H., Jarvis, E. D., Fedrigo, O., Koepfli, K.-P., Urban, L., Gemmell, N. J. & Li, H. (2022). Haplotype-resolved assembly of diploid genomes without parental data. Nature Biotechnology 40(9), 1332–1335.

Consortium, I. C. G. M. (2015). High-Resolution Linkage Map and Chromosome-Scale Genome Assembly for Cassava (Manihot esculenta Crantz) from 10 Populations. G3 Genes|Genomes|Genetics 5(1), 133–144.

Danecek, P., Bonfield, J. K., Liddle, J., Marshall, J., Ohan, V., Pollard, M. O., Whitwham, A., Keane, T., Mccarthy, S. A., Davies, R. M. & Li, H. (2021). Twelve years of SAMtools and BCFtools. Gigascience 10(2).

Doyle, J. J. & Doyle, J. L. (1987). A Rapid DNA Isolation Procedure for Small Quantities of Fresh Leaf Tissue. Phytochemical Sect., Botanical Soc. of America.

Ellinghaus, D., Kurtz, S. & Willhoeft, U. (2008). LTRharvest, an efficient and flexible software for de novo detection of LTR retrotransposons. Bmc Bioinformatics 9, 1–14.

Fu, Z.-Y., Song, J.-C. & Jameson, P. E. (2017). A rapid and cost effective protocol for plant genomic DNA isolation using regenerated silica columns in combination with CTAB extraction. Journal of Integrative Agriculture 16(8), 1682–1688.

Gabriel, L., Brůna, T., Hoff, K. J., Ebel, M., Lomsadze, A., Borodovsky, M. & Stanke, M. (2024). BRAKER3: Fully automated genome annotation using RNA-seq and protein evidence with GeneMark-ETP, AUGUSTUS, and TSEBRA. Genome Research.

Gabriel, L., Hoff, K. J., Brůna, T., Borodovsky, M. & Stanke, M. (2021). TSEBRA: transcript selector for BRAKER. Bmc Bioinformatics 22(1), 566.

Gotoh, O., Morita, M. & Nelson, D. R. (2014). Assessment and refinement of eukaryotic gene structure prediction with gene-structure-aware multiple protein sequence alignment. Bmc Bioinformatics 15, 1–13.

Gu, Z., Gu, L., Eils, R., Schlesner, M. & Brors, B. (2014). “ Circlize” implements and enhances circular visualization in R.

Han, F., Qu, Y., Chen, Y., Xu, L. A. & Bi, C. (2022). Assembly and comparative analysis of the complete mitochondrial genome of Salix wilsonii using PacBio HiFi sequencing. Front Plant Sci 13, 1031769.

Iwata, H. & Gotoh, O. (2012). Benchmarking spliced alignment programs including Spaln2, an extended version of Spaln that incorporates additional species-specific features. Nucleic Acids Research 40(20), e1611-e161.

Jayakodi, M., Padmarasu, S., Haberer, G., Bonthala, V. S., Gundlach, H., Monat, C., Lux, T., Kamal, N., Lang, D., Himmelbach, A., Ens, J., Zhang, X. Q., Angessa, T. T., Zhou, G., Tan, C., Hill, C., Wang, P., Schreiber, M., Boston, L. B., Plott, C., Jenkins, J., Guo, Y., Fiebig, A., Budak, H., Xu, D., Zhang, J., Wang, C., Grimwood, J., Schmutz, J., Guo, G., Zhang, G., Mochida, K., Hirayama, T., Sato, K., Chalmers, K. J., Langridge, P., Waugh, R., Pozniak, C. J., Scholz, U., Mayer, K. F. X., Spannagl, M., Li, C., Mascher, M. & Stein, N. (2020). The barley pan-genome reveals the hidden legacy of mutation breeding. Nature 588(7837), 284–289.

Kim, D., Langmead, B. & Salzberg, S. L. (2015). HISAT: a fast spliced aligner with low memory requirements. Nature Methods 12(4), 357–360.

Kim, D., Paggi, J. M., Park, C., Bennett, C. & Salzberg, S. L. (2019). Graph-based genome alignment and genotyping with HISAT2 and HISAT-genotype. Nature Biotechnology 37(8), 907–915.

Kolmogorov, M., Yuan, J., Lin, Y. & Pevzner, P. A. (2019). Assembly of long, error-prone reads using repeat graphs. Nature Biotechnology 37(5), 540–546.

Kovaka, S., Zimin, A. V., Pertea, G. M., Razaghi, R., Salzberg, S. L. & Pertea, M. (2019). Transcriptome assembly from long-read RNA-seq alignments with StringTie2. Genome Biology 20, 1–13.

Kuon, J.-E., Qi, W., Schläpfer, P., Hirsch-Hoffmann, M., Von Bieberstein, P. R., Patrignani, A., Poveda, L., Grob, S., Keller, M. & Shimizu-Inatsugi, R. (2019). Haplotype-resolved genomes of geminivirus-resistant and geminivirus-susceptible African cassava cultivars. BMC biology 17, 1–15.

Kuznetsov, D., Tegenfeldt, F., Manni, M., Seppey, M., Berkeley, M., Kriventseva, Evgenia V. & Zdobnov, E. M. (2022). OrthoDB v11: annotation of orthologs in the widest sampling of organismal diversity. Nucleic Acids Research 51(D1), D445-D451.

Landi, M., Shah, T., Falquet, L., Niazi, A., Stavolone, L., Bongcam-Rudloff, E. & Gisel, A. (2023). Haplotype-resolved genome of heterozygous African cassava cultivar TMEB117 (Manihot esculenta). Scientific Data 10(1), 887.

Li, H. (2018). Minimap2: pairwise alignment for nucleotide sequences. Bioinformatics 34(18), 3094–3100.

Li, H., Handsaker, B., Wysoker, A., Fennell, T., Ruan, J., Homer, N., Marth, G., Abecasis, G., Durbin, R. & Subgroup, G. P. D. P. (2009). The sequence alignment/map format and SAMtools. Bioinformatics 25(16), 2078–2079.

Lomsadze, A., Ter-Hovhannisyan, V., Chernoff, Y. O. & Borodovsky, M. (2005). Gene identification in novel eukaryotic genomes by self-training algorithm. Nucleic Acids Research 33(20), 6494–6506.

Lozano, R., Hamblin, M. T., Prochnik, S. & Jannink, J. L. (2015). Identification and distribution of the NBS-LRR gene family in the Cassava genome. Bmc Genomics 16.

Manni, M., Berkeley, M. R., Seppey, M. & Zdobnov, E. M. (2021). BUSCO: assessing genomic data quality and beyond. Current Protocols 1(12), e323.

Mansfeld, B. N., Boyher, A., Berry, J. C., Wilson, M., Ou, S., Polydore, S., Michael, T. P., Fahlgren, N. & Bart, R. S. (2021). Large structural variations in the haplotype-resolved African cassava genome. The Plant Journal 108(6), 1830–1848.

Meckoni, S. N., Nass, B. & Pucker, B. (2023). Phylogenetic placement of Ceratophyllum submersum based on a complete plastome sequence derived from nanopore long read sequencing data. BMC Res Notes 16(1), 187.

Ou, S. & Jiang, N. (2018). LTR_retriever: a highly accurate and sensitive program for identification of long terminal repeat retrotransposons. Plant Physiology 176(2), 1410-1422.

Ou, S. & Jiang, N. (2019). LTR_FINDER_parallel: parallelization of LTR_FINDER enabling rapid identification of long terminal repeat retrotransposons. Mobile DNA 10(1), 48.

Ou, S., Su, W., Liao, Y., Chougule, K., Agda, J. R., Hellinga, A. J., Lugo, C. S. B., Elliott, T. A., Ware, D. & Peterson, T. (2019). Benchmarking transposable element annotation methods for creation of a streamlined, comprehensive pipeline. Genome Biology 20, 1–18.

Pertea, G. & Pertea, M. (2020). GFF utilities: GffRead and GffCompare. F1000Research 9.

Pertea, M., Kim, D., Pertea, G. M., Leek, J. T. & Salzberg, S. L. (2016). Transcript-level expression analysis of RNA-seq experiments with HISAT, StringTie and Ballgown. Nat Protoc 11(9), 1650–1667.

Prochnik, S., Marri, P. R., Desany, B., Rabinowicz, P. D., Kodira, C., Mohiuddin, M., Rodriguez, F., Fauquet, C., Tohme, J., Harkins, T., Rokhsar, D. S. & Rounsley, S. (2012). The Cassava Genome: Current Progress, Future Directions. Trop Plant Biol 5(1), 88–94.

Pucker, B., Holtgräwe, D., Rosleff Sörensen, T., Stracke, R., Viehöver, P. & Weisshaar, B. (2016). A De Novo Genome Sequence Assembly of the Arabidopsis thaliana Accession Niederzenz-1 Displays Presence/Absence Variation and Strong Synteny. Plos One 11(10), e0164321.

Qi, W., Lim, Y. W., Patrignani, A., Schlapfer, P., Bratus-Neuenschwander, A., Gruter, S., Chanez, C., Rodde, N., Prat, E., Vautrin, S., Fustier, M. A., Pratas, D., Schlapbach, R. & Gruissem, W. (2022). The haplotype-resolved chromosome pairs of a heterozygous diploid African cassava cultivar reveal novel pan-genome and allele-specific transcriptome features. Gigascience 11.

Quinlan, A. R. (2014). BEDTools: the Swiss-army tool for genome feature analysis. Current protocols in bioinformatics 47(1), 11.12. 1-11.12. 34.

Quinlan, A. R. & Hall, I. M. (2010). BEDTools: a flexible suite of utilities for comparing genomic features. Bioinformatics 26(6), 841–842.

Ramu, P., Esuma, W., Kawuki, R., Rabbi, I. Y., Egesi-, C., Bredeson, J. V., Bart, R. S., Verma, J., Buckler, E. S. & Lu, F. (2017). Cassava haplotype map highlights fixation of deleterious mutations during clonal propagation. Nature Genetics 49(6), 959-+.

Shafin, K., Pesout, T., Lorig-Roach, R., Haukness, M., Olsen, H. E., Bosworth, C., Armstrong, J., Tigyi, K., Maurer, N., Koren, S., Sedlazeck, F. J., Marschall, T., Mayes, S., Costa, V., Zook, J. M., Liu, K. J., Kilburn, D., Sorensen, M., Munson, K. M., Vollger, M. R., Monlong, J., Garrison, E., Eichler, E. E., Salama, S., Haussler, D., Green, R. E., Akeson, M., Phillippy, A., Miga, K. H., Carnevali, P., Jain, M. & Paten, B. (2020). Nanopore sequencing and the Shasta toolkit enable efficient de novo assembly of eleven human genomes. Nature Biotechnology 38(9), 1044–1053.

Sheat, S., Fuerholzner, B., Stein, B. & Winter, S. (2019). Resistance Against Cassava Brown Streak Viruses From Africa in Cassava Germplasm From South America. Front Plant Sci 10.

Sheat, S., Margaria, P. & Winter, S. (2021). Differential Tropism in Roots and Shoots of Resistant and Susceptible Cassava (Manihot esculenta Crantz) Infected by Cassava Brown Streak Viruses. Cells 10(5).

Sheat, S. & Winter, S. (2023). Developing broad-spectrum resistance in cassava against viruses causing the cassava mosaic and the cassava brown streak diseases. Front Plant Sci 14, 1042701.

Shi, J. & Liang, C. (2019). Generic Repeat Finder: A High-Sensitivity Tool for Genome-Wide De Novo Repeat Detection. Plant Physiology 180(4), 1803–1815.

Stanke, M., Diekhans, M., Baertsch, R. & Haussler, D. (2008). Using native and syntenically mapped cDNA alignments to improve de novo gene finding. Bioinformatics 24(5), 637-644.

Stanke, M., Schöffmann, O., Morgenstern, B. & Waack, S. (2006). Gene prediction in eukaryotes with a generalized hidden Markov model that uses hints from external sources. Bmc Bioinformatics 7(1), 62.

Su, W., Gu, X. & Peterson, T. (2019). TIR-Learner, a new ensemble method for TIR transposable element annotation, provides evidence for abundant new transposable elements in the maize genome. Molecular Plant 12(3), 447–460.

Sun, Y., Shang, L., Zhu, Q.-H., Fan, L. & Guo, L. (2022). Twenty years of plant genome sequencing: achievements and challenges. Trends in Plant Science 27(4), 391–401.

Tang, H., Zhang, X., Miao, C., Zhang, J., Ming, R., Schnable, J. C., Schnable, P. S., Lyons, E. & Lu, J. (2015). ALLMAPS: robust scaffold ordering based on multiple maps. Genome Biology 16, 1–15.

Wang, M., Zhang, L., Tong, S., Jiang, D. & Fu, Z. (2022). Chromosome-level genome assembly of a xerophytic plant, Haloxylon ammodendron. DNA Research 29(2).

Xiong, W., He, L., Lai, J., Dooner, H. K. & Du, C. (2014). HelitronScanner uncovers a large overlooked cache of Helitron transposons in many plant genomes. Proceedings of the National Academy of Sciences 111(28), 10263–10268.

Xu, X.-D., Zhao, R.-P., Xiao, L., Lu, L., Gao, M., Luo, Y.-H., Zhou, Z.-W., Ye, S.-Y., Qian, Y.-Q. & Fan, B.-L. (2023). Telomere-to-telomere assembly of cassava genome reveals the evolution of cassava and divergence of allelic expression. Horticulture Research 10(11), uhad200.

Xu, Z. & Wang, H. (2007). LTR_FINDER: an efficient tool for the prediction of full-length LTR retrotransposons. Nucleic Acids Research 35(Suppl_2), W265–W268.

Zhang, R.-G., Li, G.-Y., Wang, X.-L., Dainat, J., Wang, Z.-X., Ou, S. & Ma, Y. (2022). TEsorter: An accurate and fast method to classify LTR-retrotransposons in plant genomes. Horticulture Research 9.

Zhang, Y., Park, C., Bennett, C., Thornton, M. & Kim, D. (2021). Rapid and accurate alignment of nucleotide conversion sequencing reads with HISAT-3N. Genome Research 31(7), 1290–1295.

